# Epithelial and γδ T cell CD47 have complementary yet distinct roles in regulating γδ intraepithelial lymphocyte migration

**DOI:** 10.64898/2026.08.19.745759

**Authors:** Ananya Parthasarathy, Matthew A. Fischer, Charles A. Parkos, Karen L. Edelblum

## Abstract

Intraepithelial lymphocytes expressing the γδ T cell receptor (γδ IEL) continuously survey the intestinal epithelium to promote mucosal host defense. Although γδ IELs migrate in and out of the lateral intercellular space (LIS) between adjacent enterocytes, the molecular mechanisms governing their migratory behavior are incompletely understood. Based on the known role of CD47, or integrin associated protein (IAP), in mediating neutrophil transepithelial migration, we investigated whether CD47 expression reflects a conserved mechanism regulating γδ IEL surveillance behavior. Here, we report that conditional CD47 deletion on intestinal epithelial cells or γδ T cells had no effect on IEL composition. Using intravital imaging, we identified complementary roles for CD47 on γδ IELs and epithelial cells, with epithelial CD47 restricting γδ IEL motility and γδ T-cell-derived CD47 promoting cell migration. Further investigation revealed that both CD47 and CD18 contribute to γδ IEL surveillance behavior, although CD47 regulates γδ IEL migration in a CD18-independent manner.

## Introduction

The intestinal barrier is made up of a single layer of epithelial cells to separate the mucosal immune system from microbes in the gut lumen. Within the epithelium are a specialized subset of T cells, known as intraepithelial lymphocytes (IEL) that serve as a first line of defense against invading pathogens^1^. IELs are broadly classified into two subsets, induced and natural IELs^2^. Induced IELs are conventional TCRαβ cells expressing either CD8αβ or CD4, whereas natural IELs express CD8αα and either TCRαβ or TCRγδ. These natural IELs are considered unconventional since their activation can be MHC-independent. In the murine small intestine, γδ IELs make up about half of the IEL compartment and continuously patrol the epithelium by migrating along the basement membrane and within the lateral intercellular space (LIS) between neighboring enterocytes^3^. Complementing their innate-like functions^4^, γδ IEL surveillance behavior is essential to limit the translocation of luminal microbes and enteric pathogens across the epithelial barrier^3,5^.

As γδ IELs survey the epithelium, these lymphocytes migrate from the basolateral aspect of the epithelium into the LIS toward the tight junction, where they transiently remain before reversing direction and exiting the monolayer^3^. These sentinels do not cross the tight junction, unlike neutrophils that migrate through the apical junctional complex into the lumen under inflammatory conditions^3,6^. The series of molecular interactions governing neutrophil transepithelial migration (TEM) is relatively well-characterized^6^ and, in part, relies on the expression of CD47 or integrin associated protein (IAP), a ubiquitously expressed cell surface protein characterized by a five-transmembrane domain connected to an extracellular IgV loop by disulfide bonds^7–9^. CD47 is also considered a marker of self, or a ‘don’t-eat-me’ signal, which prevents phagocytosis via recognition signal regulatory protein α (SIRPα)^10^. While it has been reported that CD47-mediated interactions regulate neutrophil TEM^11,12^, more recent studies strongly implicate CD47 as a mediator of CD11b/CD18 (α_M_β_2_ integrin) function^9^, consistent with its reported role in regulating other integrin functions^10,13^. Based on the migratory behavior of neutrophils, we hypothesized that CD47 expression may reflect a conserved mechanism by which leukocytes, including γδ IELs, migrate into the LIS.

We previously demonstrated a role for CD103 (α_E_β_7_) in γδ IEL motility^3^, yet the role of other integrins in modulating γδ IEL surveillance behavior remains unclear. Notably, γδ IELs express CD47 along with multiple integrins, including CD11a (α_L_), CD11c (α_X_), and CD18 (β_2_)^14^. In this study, we assessed the respective contribution of epithelial– and γδ T-cell-derived CD47 in regulating γδ IEL motility. We report that CD47 deletion on either cell type had no effect on IEL composition or frequency under homeostatic conditions, yet loss of γδ T cell CD47 led to a reduction in CD11c/CD18^+^ γδ IELs. Using intravital imaging, we identified complementary roles for CD47 on γδ IELs and epithelial cells, with epithelial CD47 restricting γδ IEL motility and γδ T-cell-derived CD47 promoting cell migration. Based on the role of neutrophil CD47 in regulating the expression of integrins in *cis*, we investigated the relationship between CD47 and CD11c/CD18 in γδ IELs. We found that loss of CD18 binding results in impaired γδ IEL motility; however, disruption of CD11c or CD18 ligation results in enhanced migration in CD47-deficient, but not wildtype γδ IELs. Although loss of epithelial CD47 enhanced γδ IEL migratory speed, this was not sufficient to confer γδ IEL-mediated protection against acute pathogen translocation, thus indicating that increased migration into the LIS is the rate-limiting step in preventing microbial invasion. Taken together, we report that both CD47 and CD18 contribute to γδ IEL surveillance behavior; however, CD47 regulates γδ IEL motility in a CD11c/CD18-independent manner.

## Materials and Methods

### Animals

All experiments were performed on 7-15-week-old C57BL/6 mice of both sexes maintained under specific pathogen-free conditions including the absence of murine norovirus and *Helicobacter* spp. Mice were housed in Allentown caging with aspen shavings, autoclaved 5010 chow and RO water, with a standard 12-hour light-dark cycle. To obtain MNV-free mice, TcrdH2BEGFP (TcrdEGFP) mice were rederived from Jackson Laboratory (RRID: IMSR_JAX:016941) and are denoted as wildtype (WT) in our studies. TcrdEGFP reporter mice were crossed to global CD47 KO mice (RRID: IMSR_JAX:003173) or intestinal epithelial cell-specific CD47 KO (CD47 KO^IEC^) mice that were generated by crossing CD47^flx^ mice to those expressing a constitutive villin-Cre as previously described^15^. Littermates not expressing Cre were used as controls (CD47 WT^IEC^). CD47^flx^ mice were crossed to those expressing an inducible TcrdCreER, provided by Dr. Yuan Zhuang (Duke University)(RRID: IMSR_JAX:031679) and *Rosa26*-tdTomato-LSL mice purchased from Jackson Laboratory (RRID: IMSR_JAX:007914) to generate tdTomato^+^ γδ T-cell-specific CD47 KO (CD47 KO^γδ^) mice. CD47 KO^γδ^ mice or R26^tdTom^; CD47^flx/+^; TcrdCreER littermate controls were treated daily with tamoxifen (2 mg, i.p., MedChemExpress) for 5 days and experiments performed after 2 weeks to allow sufficient Cre recombination of the floxed alleles. To investigate the role of CD11c or CD18 in CD47-mediated γδ IEL migration, CD47 het^γδ^ and CD47 KO^γδ^ mice were treated with 20 μg (i.p.) of αCD11c or 200 μg αCD18 (BioXCell, BE0038 or BE0009, respectively) for 1 h prior to imaging. All studies were conducted in an Association of the Assessment and Accreditation of Laboratory Animal Care-accredited (IACUC) facility according to protocols approved by the Center for Comparative Medicine and Surgery at the Icahn School of Medicine at Mount Sinai.

### IEL and epithelial cell isolation

Murine small intestinal IELs were isolated as previously described^5^. Briefly, the small intestine was excised, opened longitudinally following the removal of Peyer’s patches, cut into one cm pieces and incubated shaking in Hank’s balanced salt solution (HBSS)(Sigma) containing 3 mM EDTA and 7.5% FBS (ThermoFisher Scientific) at 37°C. The supernatant was collected and replaced every 20 min for 1 h after which IELs were enriched using a glass wool column (ThermoFisher Scientific) and subsequently purified using a 20/45/70% Percoll (Cytiva) density gradient. EpCAM^+^ epithelial cells isolated from this fraction were also analyzed.

### Flow cytometry

Freshly isolated cells were stained with fixed viability dye eFluor 780 and CD11b (M1/70) from eBioscience, TCRβ (H57-597), TCRδ (GL3), CD4 (RM4-5), CD47 (miap301), CD103 (2E7), CD326/EpCAM (G8.8) from Biolegend, biotinylated CD29 (HMb1-1) from Invitrogen, CD3ε (145-2C11), CD8α (53-6.7), CD8β (YTS156.7.7), Vγ1 (2.11), Vγ4 (UC3-10A6), CD11a (M17/4), CD11c (N418), CD18 (GAME-46), and CD61 (2C9.G2) from BD Biosciences. Biotinylated Vγ7 (GL7) was provided by Rebecca O’Brien (National Jewish Health, Denver, CO). Biotinylated antibodies were subsequently stained with streptavidin-BUV737 (BD Biosciences). When necessary, cells were fixed using Cytofix/Cytoperm (Becton Dickinson). Flow cytometry was performed using a FACSymphony A5 SE (BD Biosciences) or Aurora (Cytek Biosciences) analyzers and the data was analyzed using FlowJo (version 10.10.0).

### Immunostaining

Jejunal tissue sections were fixed in 4% paraformaldehyde at room temperature for 2 hours, free aldehydes were quenched with 50 mM NH_4_Cl, followed by incubation in 30% sucrose overnight at 4°C. The buffered tissue was subsequently embedded in optimal cutting temperature medium (VWR) and stored at –20°C. To quantify γδ IELs, 5-7μm sections were permeabilized in 0.5% NP-40, blocked in 10% normal goat serum (NGS) and stained with anti-laminin (Sigma-Aldrich, L9393) and anti-rabbit AlexaFluor488 or 594 (Life Technologies, A11008 and A11012, respectively). F-actin and nuclei were stained with AlexaFluor647-conjugated phalloidin (Life Technologies A22287) and Hoechst 33342 (Life Technologies H3570). To visualize CD47 *in situ*, 5-7μm sections were blocked and permeabilized in 3% BSA and 0.5% Triton X-100 in PBS and stained with anti-CD47 (R&D Systems, AF1866) followed by anti-goat AlexaFluor647 (Life Technologies A21447) and Hoechst 33342. All slides were mounted with Prolong Glass AntiFade (ThermoFisher Scientific, P36980). Imaging was performed on an Andor Dragonfly 620 Spinning Disk confocal microscope using a Sona 4.2 sCMOS camera, PlanApo 20x/0.8 λD or PlanApo 40x/1.25 Sil λS objective and Fusion acquisition software (v. 2.4.0.14). The number of γδ IELs per 0.1 mm^2^ of jejunum was quantified using FIJI (NIH) blinded to the experimental condition/genotype.

### Intravital microscopy

Intravital imaging was performed as previously described^16^. Briefly, each mouse was anesthetized and a retroorbital injection of Hoechst 33342 was administered to label nuclei. A 2-3 cm long region of jejunum was exposed, cauterized, and opened to reveal the gut lumen. The mouse was then placed onto a glass bottom 35 mm dish, the exposed mucosa bathed in HBSS with AlexaFluor633 hydrazide (Invitrogen, A30634) to visualize the intestinal lumen. Timelapse microscopy was performed on an Andor Dragonfly 620 Spinning Disk confocal microscope equipped with an iXon Life 888 EMCCD camera, PlanApo 40x/1.25 Sil λS objective and Fusion software (v. 2.4.0.14). 15 μm z-stacks (1.5 μm slice) were acquired every 20 s for 30 min after which 4D reconstructions of the time-lapse videos were rendered using Imaris (v. 10.1.1) and an autoregressive tracking algorithm was employed to quantify γδ IEL migration. Arrest coefficients were calculated as the proportion of the track duration in which the instantaneous speed of a γδ IEL was less than 2 μm/min.

### Salmonella infection

DsRed-labeled *Salmonella* Typhimurium (strain SL3201) was provided by Dr. Andrew Neish (Emory University). After culturing *S.* Typhimurium in Luria-Bertani medium containing 100 mM ampicillin, the jejunal mucosa was infected by anesthetizing mice and performing laparotomies as previously described above^5,16^. 10^8^ CFU DsRed-SL3201 was applied to the exposed mucosa for 30 min.

### Statistical analysis

All statistical analyses were performed using GraphPad Prism (v. 10.4.1). For normally distributed data comparing two variables, unpaired t-tests were performed, whereas Mann Whitney U tests were used for non-parametric data. For comparisons across three or more variables, one-way and two-way ANOVAs with the appropriate post-hoc tests were used to compare normally distributed data, and Kruskal-Wallis used for non-parametric data sets. All experiments include at least two independent experiments with a minimum of n=3 mice per genotype/condition.

## Results

### CD47 is not required for maintenance of the γδ IEL compartment

Based on the ubiquitous expression of CD47, we generated CD47 conditional knockout mice to elucidate the specific contribution of epithelial– and γδ T-cell-derived CD47 to γδ IEL migration. To this end, CD47^flx^; vilCre (CD47 KO^IEC^) mice were crossed to a GFP γδ T cell reporter strain to allow visualization of γδ T cells in the absence of intestinal epithelial CD47 expression (Fig. 1A). Loss of epithelial CD47 was confirmed by immunostaining (Fig. 1B,C); however, we were unable to visualize CD47 on the IEL cell surface, which may be due to the comparatively low level of CD47 expression on these cells. Reciprocally, CD47^flx^ mice were crossed to those expressing a *Rosa26*-tdTomato reporter and an inducible Cre driven downstream of the *Tcrd* promoter (TcrdCreER)(CD47 KO^γδ^ mice), such that treatment with tamoxifen generates tdTom^+^ CD47-deficient γδ T cells (Fig. 1D).

**Figure 1.**
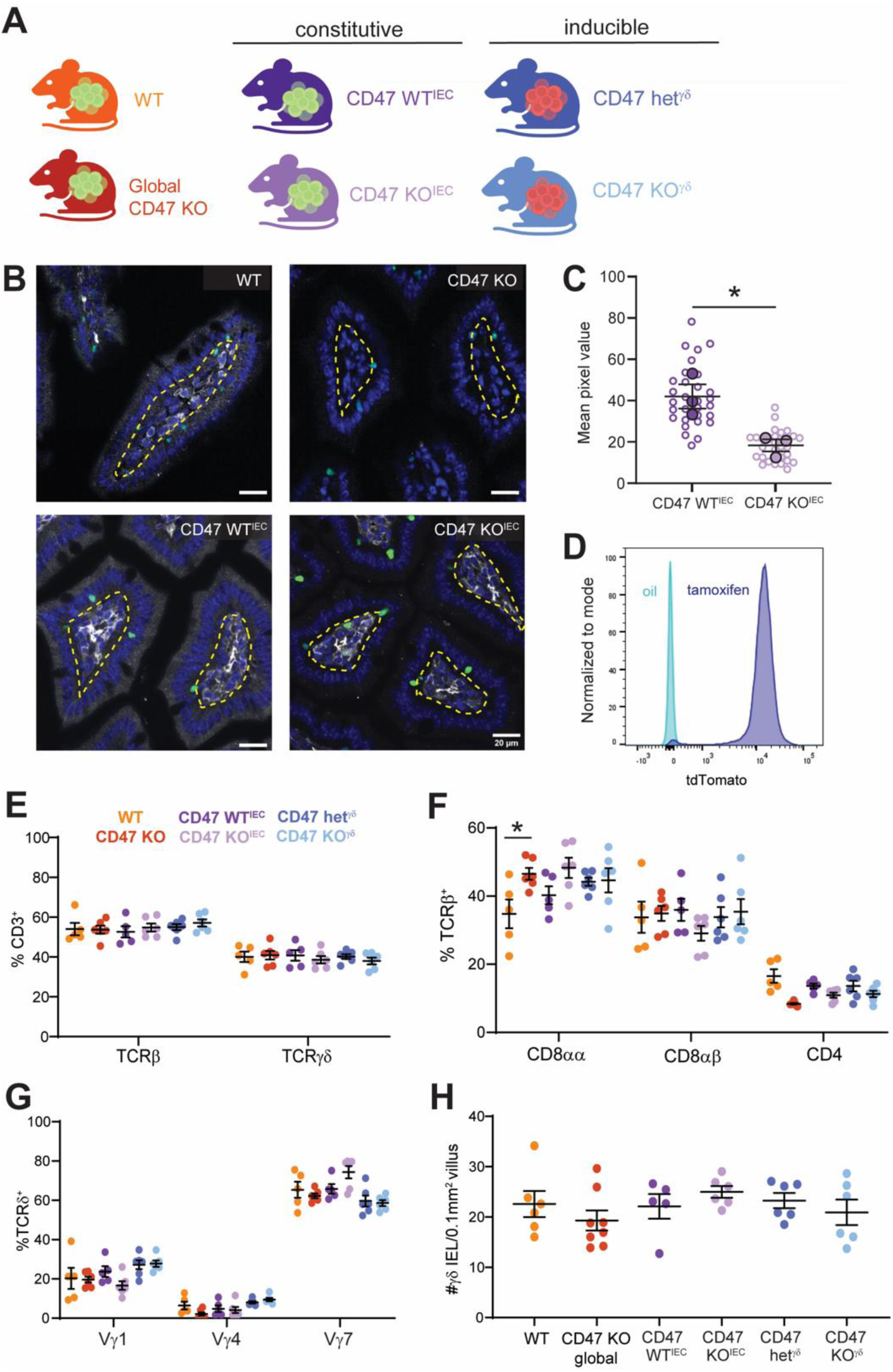
Conditional deletion of CD47 has no effect on γδ IEL number or overall IEL composition. (A) Schematic of mouse strains used to assess the cell-specific contribution of CD47 on γδ IEL phenotype and migration made with Biorender (B) Immunostaining for CD47 (white) in jejunum of WT, global CD47 KO, CD47 WT^IEC^ or CD47 KO^IEC^ mice. γδ IELs are shown in green, nuclei in blue, and a yellow dashed line approximates the basement membrane. Scale bar = 20μm. (C) Morphometric analysis of the fluorescence intensity of CD47 in intestinal epithelium. Approximately 60-75 villi were analyzed (open circles), with the mean pixel value per mouse represented by filled circles. n=3 mice. (D) Representative histogram showing R26^tdTomato^ expression in oil– or tamoxifen-treated CD47 KO^γδ^ mice. Flow cytometric analysis indicating (E) the frequency of TCRβ and TCRγδ gated on CD3, (F) CD4, CD8αα, or CD8αβ gated on TCRβ or (G) Vγ IEL subsets isolated from WT, global CD47 KO or conditional CD47 KO mice. n=5-6 mice. (H) Morphometric analysis of jejunal γδ IEL number from global and conditional CD47 knockout mice compared to WT and littermate controls. n=5-8 mice. All data are shown as mean ± SEM from at least 2 independent experiments. Each data point represents an individual mouse unless otherwise noted. Statistical analysis: (C) unpaired *t*-test, (E-G) two-way or (H) one-way ANOVA with Sidak’s posthoc test. *P<0.05

Based on the reported role of CD47 as a marker of “self” to prevent phagocytosis by macrophages^17,18^, we first asked whether conditional loss of CD47 alters the composition of the IEL compartment. Neither deletion of epithelial, nor γδ T cell CD47 significantly affected the relative proportion of induced or natural IEL subsets within the small intestine. However, we observed a minor increase in CD8αα TCRαβ IELs in global CD47 KO compared to WT controls that was not recapitulated in the conditional CD47 KO lines (Fig. 1E,F, Fig. S1). Further, the frequency of individual Vγ IEL subsets remained unchanged following deletion of CD47 in either cell type (Fig. 1G). Morphometric analysis of GFP^+^ or tdTom^+^ γδ IELs in the respective conditional knockout strains revealed no difference in total γδ IEL number relative to littermate controls (Fig. 1H). Together, these data indicate that CD47 expression by either cell type is dispensable for the development and maintenance of the IEL compartment.

### Epithelial CD47 promotes γδ IEL retention within the LIS

Previous studies demonstrated a critical role for CD47 in neutrophil TEM through interaction with multiple ligands including signal-regulatory protein alpha (SIRPα), thrombospondin-1 (TSP-1) and CD18^7–9,11^. Given that γδ IELs do not express SIRPα or TSP-1^19^, we hypothesized that epithelial CD47 may similarly promote γδ IEL migration into the LIS. To test this, we visualized γδ IEL migration within the jejunal mucosa of CD47 KO^IEC^ mice and wildtype (CD47 WT^IEC^) littermate controls using intravital microscopy. Loss of epithelial CD47 led to an increase in γδ IEL average track speed, which was associated with reduced dwell time of γδ IELs within the LIS (Fig. 2A,B). Despite this, the overall frequency of γδ IELs within the LIS was similar between CD47 KO^IEC^ and CD47 WT^IEC^ mice (Fig. 2C), indicating that epithelial CD47 does not regulate the incidence of γδ IEL entry into and exit from the epithelium. Further, we observed no difference in the number of times an individual enterocyte is contacted by γδ IEL between the two genotypes (Fig. 2D), demonstrating that γδ IEL surveillance behavior is not compromised in the absence of epithelial CD47 expression. These findings suggest that epithelial CD47 may function as a ‘stop’ signal to promote the retention of γδ IELs within the LIS with the loss of this retention signal potentially reflecting a faster off-rate of CD47-integrin binding resulting an increase in overall migratory speed.

**Figure 2.**
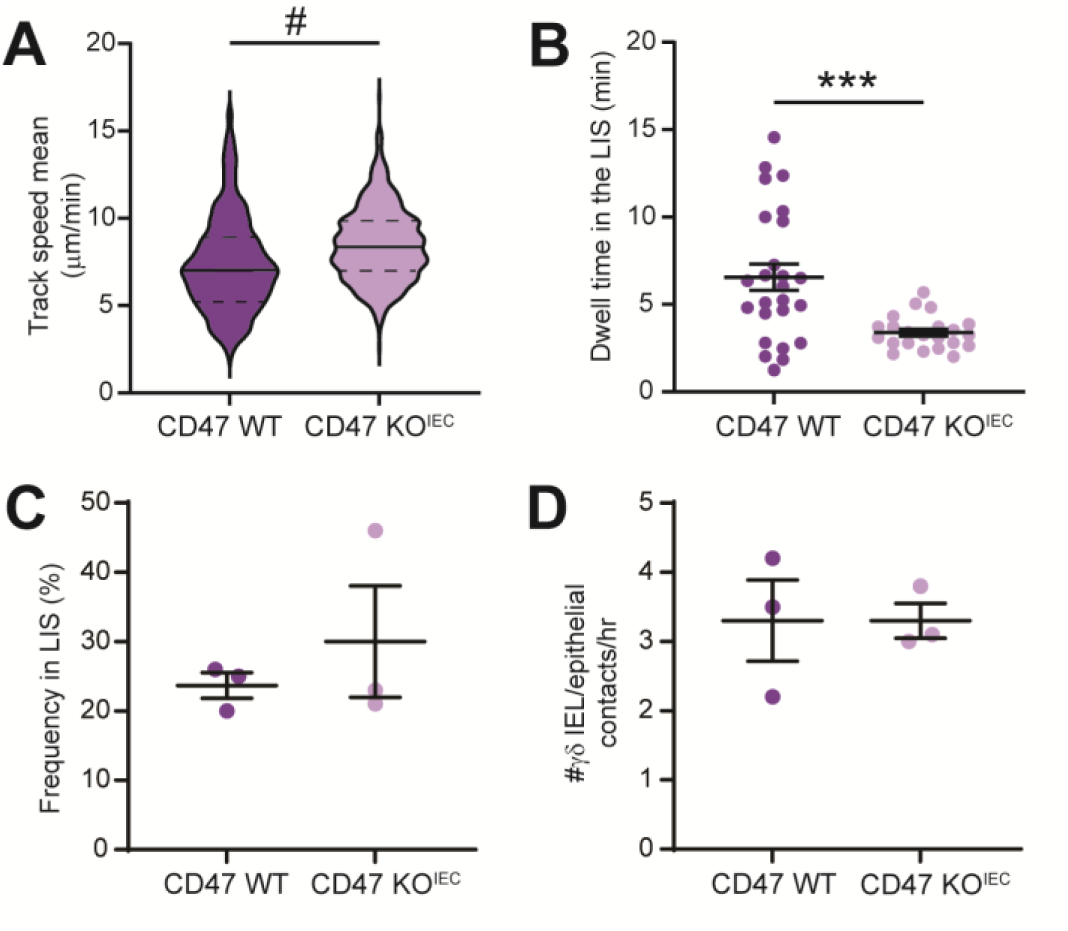
Epithelial CD47 limits γδ IEL motility and facilitates arrest within the LIS. Intravital imaging of jejunal mucosa was performed on TcrdEGFP; CD47 WT^IEC^ or CD47 KO^IEC^ mice. (A) γδ IEL mean track speed, (B) dwell time of individual γδ IELs within the LIS, (C) the frequency of all γδ IELs localized within the LIS and (D) the number of times an individual epithelial cell interacts with an γδ IEL over the course of an hour is shown. (A) Median and quartiles are shown. (B-D) Data represent the mean ± SEM from 3 mice of each genotype. (B) Each data point represents an individual γδ IEL, (C,D) each data point represents an individual mouse. n=438 and 559 tracks for CD47 WT^IEC^ and CD47 KO^IEC^ respectively. Statistical analysis: (A) Mann-Whitney U test; (B-D) unpaired t-test. ***P<0.001, #P<0.0001.

### γδ T cell CD47 facilitates contact with the IECs by enhancing migratory speed

To determine the contribution of γδ IEL CD47 to γδ IEL surveillance behavior, we performed live imaging of tdTom^+^ γδ IELs in the jejunal mucosa of CD47 KO^γδ^ and littermate controls heterozygous for the CD47^flx^ allele (CD47 het^γδ^). In contrast to our observations in CD47 KO^IEC^ mice, we found that the average track speed of γδ IELs was reduced in CD47 KO^γδ^ mice, coupled with an increased dwell time within the LIS (Fig. 3A,B, Supplemental Video 1). Whereas loss of γδ T cell CD47 had no effect on the frequency of γδ IELs within the LIS, the decrease in migratory speed leads to a trend toward reduced γδ IEL/epithelial interactions (Fig. 3C,D). Collectively, these data demonstrate that γδ T-cell-derived CD47 contributes to steady-state surveillance behavior by promoting γδ IEL motility and exit from the LIS.

**Figure 3.**
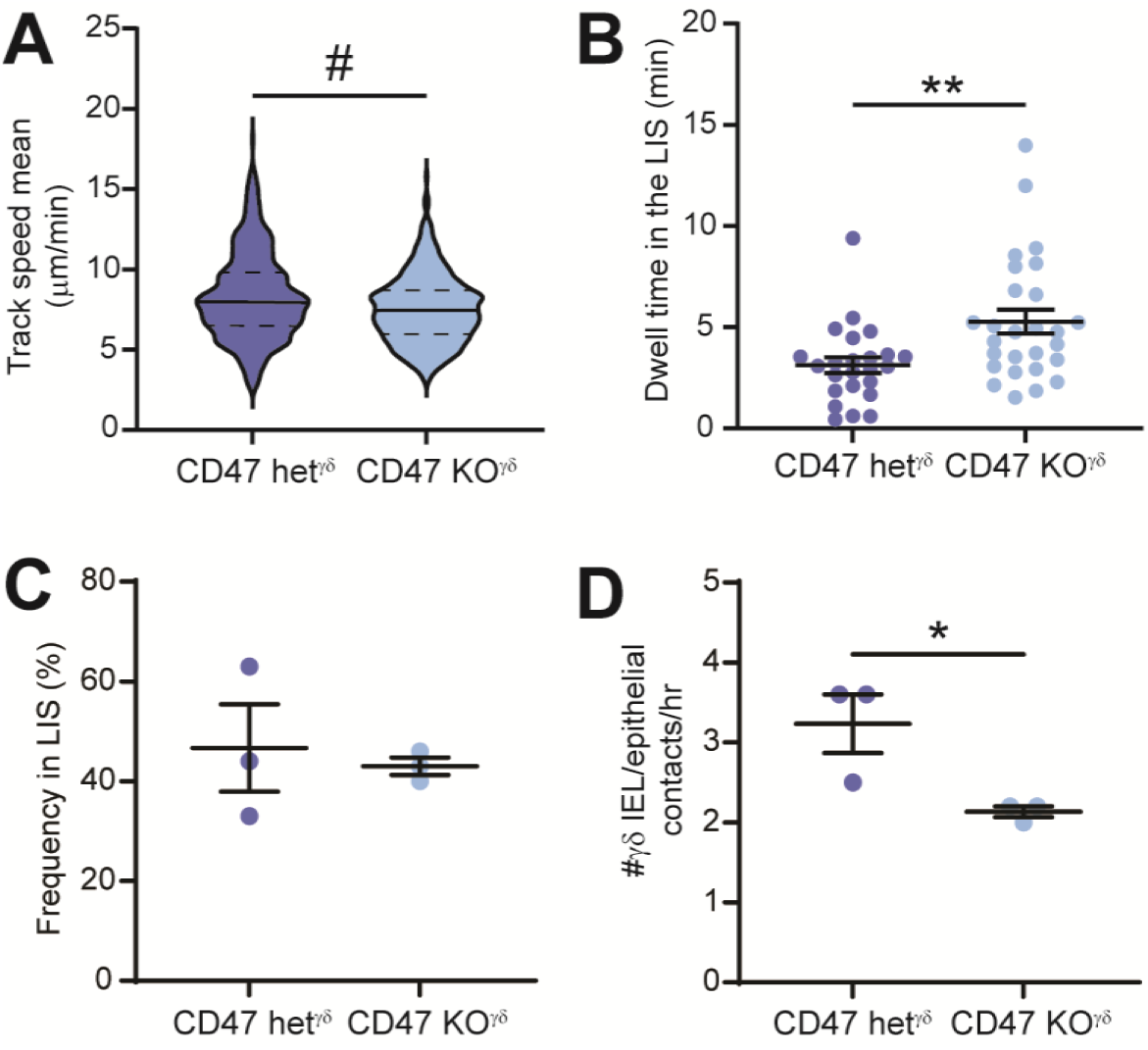
Loss of γδ T cell CD47 reduces γδ IEL motility and promotes their retention within the LIS. Intravital imaging of tdTomato^+^ γδ IELs was performed in CD47 het^γδ^ and CD47 KO^γδ^ jejunal mucosa. (A) γδ IEL mean track speed, (B) dwell time of individual γδ IELs within the LIS, (C) the frequency of all γδ IELs localized within the LIS and (D) the number of times an individual epithelial cell interacts with an γδ IEL per hour is shown. (A) Median and quartiles are shown. (B-D) Data represent the mean ± SEM from 3 mice of each genotype. (B) Each data point represents an individual γδ IEL, (C-D) each data point represents an individual mouse. n=414 and 424 tracks for CD47 het^γδ^ and CD47 KO^γδ^ respectively. Statistical analysis: (A, D) Mann-Whitney U test; (B, C) unpaired t-test. *P<0.05, **P<0.01, #P<0.0001.

These findings define complementary yet distinct roles for epithelial and γδ T cell CD47 in modulating γδ IEL motility. Since epithelial CD47 appears to act as a ‘stop’ signal to promote γδ IEL retention in the LIS, whereas our data suggests that γδ T cell CD47 serves as a ‘go’ signal, we next asked how global loss of CD47 impacts γδ IEL motility. Intravital imaging of global CD47 KO and WT mucosa revealed no substantial differences among metrics of γδ IEL migration between the two genotypes (Fig. 4). The lack of a migratory phenotype was surprising, but not entirely unexpected given the observed opposing effects of epithelial and γδ T cell CD47 on γδ IEL surveillance behavior. These results serve as a reminder that (1) the absence of a phenotype using a global knockout strategy may mask the cell-specific function of a given molecule and (2) defining the contribution of a ubiquitously expressed protein on individual cell types is essential to elucidate the molecular underpinning of cellular interactions.

**Figure 4.**
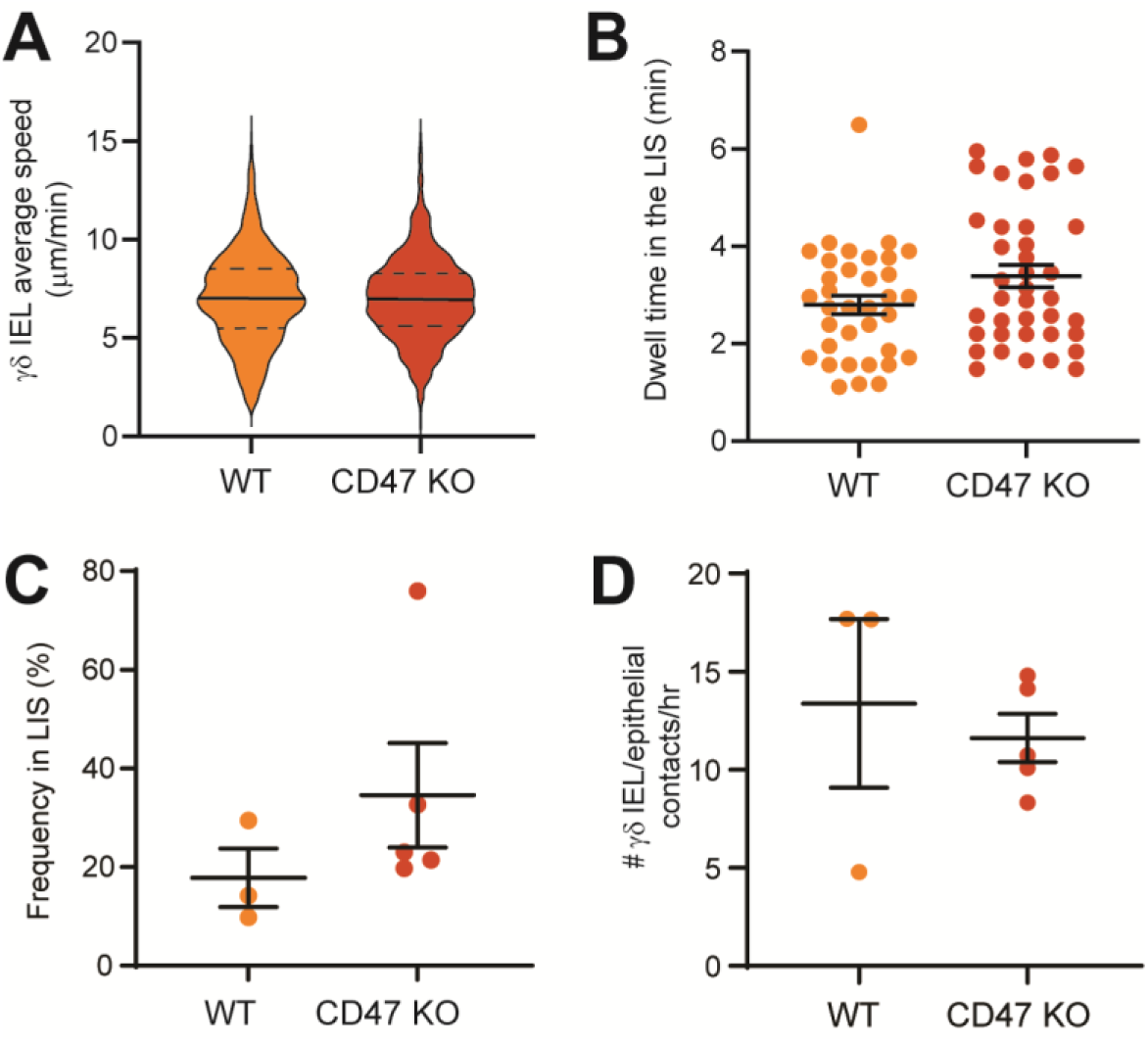
Global loss of CD47 expression has no effect on γδ IEL migratory behavior. Intravital imaging of jejunal mucosa was performed on TcrdEGFP; WT or CD47 KO mice. (A) γδ IEL mean track speed, (B) dwell time of individual γδ IELs within the LIS, (C) the frequency of all γδ IELs localized within the LIS and (D) the number of times an individual epithelial cell interacts with an γδ IEL over an hour is shown. (A) Median and quartiles are shown. (B-D) Data represent the mean ± SEM from 3-5 mice of each genotype. (B) Each data point represents an individual γδ IEL, (C-D) each data point represents an individual mouse. n=736 (WT) or 845 (CD47 KO) tracks. Statistical analysis: (A, D) Mann-Whitney U test; (B, C) student’s t-test.

### CD47 regulates γδ IEL motility in a CD11c/CD18-independent manner

The positive association of γδ T cell CD47 in promoting cell migration next led us to ask whether CD47 regulates integrin expression in *cis* within these lymphocytes. In neutrophils, CD47 upregulates the CD11b/CD18 receptor binding affinity/avidity, which facilitates binding interactions with as-of yet unidentified epithelial ligand(s) to promote neutrophil TEM^9^. Prior proteomic analyses indicate that γδ IELs express several beta integrins, including CD18 and its canonical alpha subunits, CD11a and CD11c^14^. Flow cytometric analysis of γδ IEL integrins revealed that the loss of epithelial CD47 had no effect on basal integrin expression (Fig. S2A,B), yet CD47-deficient γδ IELs exhibited a reduction in the frequency of CD11c^+^ CD18^+^ cells (Fig. 5A-C). Notably, the expression of individual integrins on the surface of integrin-positive WT and CD47-deficient γδ IELs was not altered (Fig. 5D), nor did we detect altered CD29 (β_1_ integrin) expression in CD47-deficient epithelial cells (Fig. S2C).

**Figure 5.**
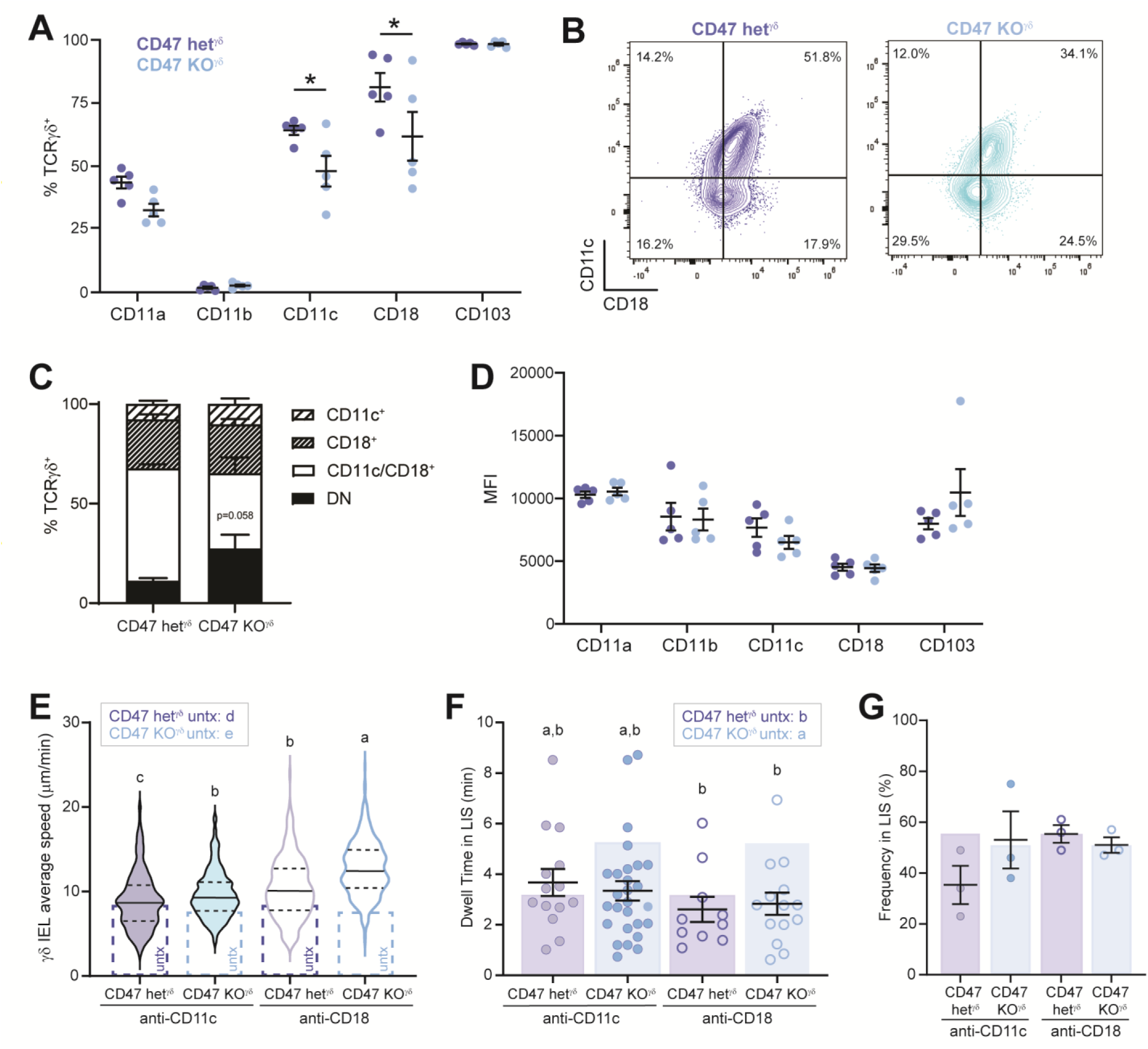
CD47 and CD18 independently modulate γδ IEL migratory behavior. (A) Frequency of CD11a, CD11b, CD11c, CD103 and CD18-expressing γδ IELs isolated from CD47 KO^γδ^ mice and CD47 het^γδ^ littermate controls. (B) Representative flow plots showing γδ IEL CD11c and CD18 expression in both genotypes. (C) Stacked bar graph highlighting the relative proportion of CD11c^+^ and/or CD18^+^ γδ IELs from data shown in (A). (D) MFI of γδ IEL integrin expression. Intravital microscopy was performed on CD47 het^γδ^ and CD47 KO^γδ^ mice pretreated with anti-CD11c or anti-CD18 (E-G). (E) Average track speed is shown. Dashed bars represent average track speed of untreated (untx) CD47 het^γδ^ and CD47 KO^γδ^ mice as shown in Fig 3A. (F) Dwell time and (G) frequency in the LIS in comparison to untx CD47 het^γδ^ and CD47 KO^γδ^ mice (shaded bars). (E) Median and quartiles are shown. (C,D,F,G) Data represent the mean ± SEM from 3-5 mice of each condition. (F) Each data point represents an individual γδ IEL, (A,D,G) each data point represents an individual mouse. n=411 CD47 het^γδ^ and 356 CD47 KO^γδ^ tracks (anti-CD11c); n=222 CD47 het^γδ^ and 328 CD47 KO^γδ^ tracks (anti-CD18). Statistical analysis: (A,C,D) two-way ANOVA with Sidak’s or (E-G) Tukey’s posthoc test. (E,F) Different letters represent statistical significance. *P<0.05, #P<0.0001.

We next asked whether blocking CD11c *in vivo* would phenocopy the migratory alterations observed in CD47 KO^γδ^ mice. Inhibiting CD11c ligation in CD47 het^γδ^ (WT) mice slightly increased γδ IEL track speed, but did not markedly impact frequency or dwell time within the LIS (Fig. 5E-G, Supplemental Video 2), indicating that CD11c blockade has a minor effect on basal γδ IEL surveillance behavior. Surprisingly, anti-CD11c exposure reversed the impaired migratory phenotype observed in CD47 KO^γδ^ mice, normalizing γδ IEL dwell time in the LIS while also promoting faster migration (Fig. 5E-G). Thus, unlike the positive association between CD47 and CD11b/CD18 in neutrophil migration, CD47 and CD11c/CD18 appear to assert differential effects on γδ IEL motility. αnti-CD18 largely phenocopied CD11c blockade, except anti-CD18 enhanced γδ IEL migratory speed to a greater extent (Supplemental Video 3). These findings indicate that another integrin besides CD11c pairs with CD18 to promote basal motility and imply that CD11c/CD18 may negatively regulate γδ IEL migration. It is unclear whether the increased dwell time of CD47-deficient γδ IELs is a result of slower migration or the absence of a CD47-mediated signal to promote exit from the LIS. Despite this, CD47-mediated modulation of γδ IEL migration within the LIS appears to occur independently of CD11c/CD18, suggesting that CD47 expression may influence an as-yet unidentified binding interaction in the LIS. Collectively, these data indicate that both CD47 and CD18 regulate γδ IEL migratory behavior, although CD47-mediated γδ IEL migration occurs through a CD11c/CD18-independent mechanism.

### Increased γδ IEL migration into the LIS confers protection against acute pathogen translocation

We previously reported that mice deficient in CD103 exhibit an increase in γδ IEL track speed, reduced retention within the LIS and enhanced frequency of γδ IELs within the LIS^5^. Further, we reported that this altered surveillance behavior confers protection against acute *Salmonella* Typhimurium invasion. Based on the similarity in the migratory phenotype between CD103-deficient mice and those lacking epithelial CD47 expression, we investigated the extent to which CD47 KO^IEC^ mice are protected from acute *Salmonella* invasion. Similar to our prior report^3^, we observed a trend of reduced *S.* Typhimurium translocation in CD103-deficient mice; however, bacterial invasion was notably increased in CD47 KO^IEC^ mice (Fig. 6A,B). CD103 KO γδ IELs migrate faster and into the LIS more frequently, whereas γδ IELs in CD47 KO^IEC^ mice simply migrate faster (Fig. 6C). Thus, the increase in frequency of γδ IELs within the LIS corresponds with improved protection against bacterial invasion, whereas ramping up γδ IEL speed and reduced dwell time in CD47 KO^IEC^ mice may compromise barrier surveillance.

**Figure 6.**
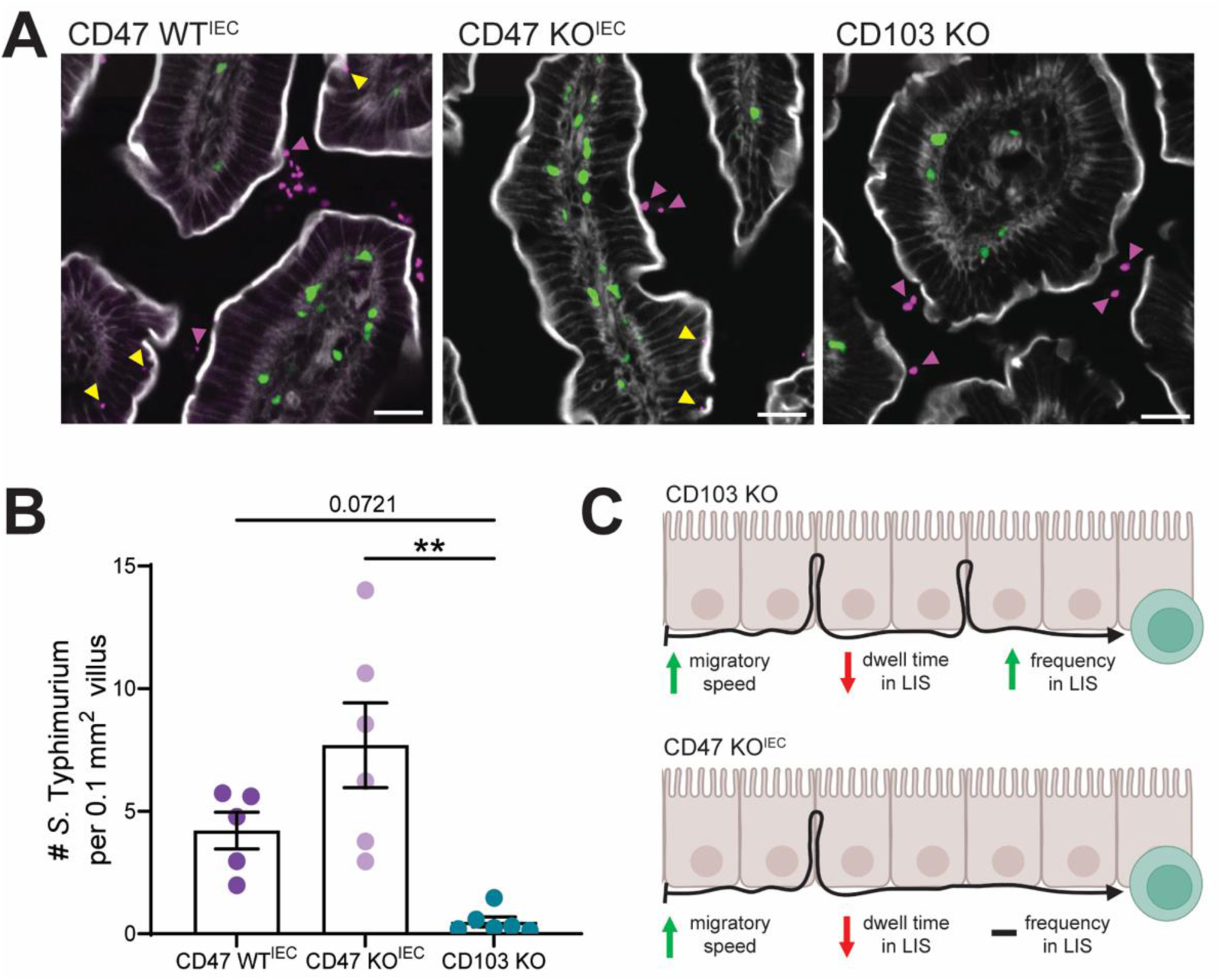
Increased frequency of γδ IELs within the LIS confers greater protection against pathogen translocation than enhanced γδ IEL migratory speed. (A) Fluorescent micrographs of *S.* Typhimurium-infected CD47 WT^IEC^, CD47 KO^IEC^ or CD103 KO mice. γδ T cells are shown in green, *S.* Typhimurium in magenta and F-actin in white. Yellow arrowheads denote bacterial translocation, whereas magenta arrowheads indicate luminal bacteria. Scale bar = 20 μm. (B) Morphometric analysis of *Salmonella* invasion at 30 min. (C) Model summarizing the role of CD103 and epithelial CD47 in γδ IEL surveillance behavior made with Biorender. n=4-6 mice. Statistical analysis: (B) one-way ANOVA with Tukey’s posthoc test. **P<0.01.

## Discussion

In this study, we identified CD47 as a key regulator of γδ IEL surveillance behavior, with epithelial and γδ T-cell-derived CD47 exhibiting distinct contributions to the migration of these sentinels along and within the epithelial barrier. We find that epithelial CD47 serves as a ‘stop’ signal to limit γδ IEL speed and facilitate retention in the LIS, whereas γδ T cell CD47 acts as a ‘go’ signal to promote motility and maintain basal surveillance. These signals appear to be complementary since γδ IEL migratory behavior is similar between WT and global CD47-deficient mice. We initially hypothesized that CD47 represents a conserved mechanism by which leukocytes navigate the intercellular space based on the requirement for neutrophil CD47 in TEM^6,9^. Whereas loss of CD47 negatively influences CD11b function in neutrophils to impair TEM^9^, we find that loss of γδ IEL CD47 reduces the frequency of CD11c/CD18^+^ γδ IELs and results in slower surveillance associated with a longer residence within the LIS. Conversely, loss of epithelial CD47 amplifies γδ IEL migratory speed, yet this is not sufficient to confer protection against acute pathogen invasion since the frequency of γδ IEL migration into the LIS remains unchanged.

CD47 has long been described as a marker of “self”, as numerous studies have reported increased phagocytosis of CD47-deficient cells^18,20^ yet the γδ IEL compartment remained unchanged in CD47 KO^γδ^ mice. Further, loss of epithelial CD47 had no measurable impact on the composition of the IEL compartment. These data are consistent with prior studies in neutrophil– or epithelial-specific CD47 KO mice in which no cell death of the respective cell population was observed^9,15^. Ablation of T cell CD47 through the use of a constitutive Lck-Cre transgene reduced peripheral T cell numbers due to cDC2-induced necroptosis in secondary lymphoid organs^21^. In our study, the use of an inducible γδ T-cell-specific Cre reduces the potential for necroptotic cell death among tissue-resident γδ IELs.

Since intestinal epithelial cells do not express the CD47 ligand SIRPα, this led us to initially hypothesize that the adhesive properties of γδ IELs may be regulated in *cis* via CD18. Given that γδ IELs express relatively little CD11b, and CD11a expression was unchanged in CD47 KO^γδ^ IELs, we focused on the role of the CD11c/CD18 heterodimer in CD47-mediated motility. CD11c is highly expressed on CD8 IEL populations and implicated in γδ IEL activation^22^, and prior work indicates that CD18 expression is dispensable for γδ IEL homeostasis^23^. Surprisingly, a clear role for either integrin in γδ IEL migration or function has yet to be determined. Use of a pan-CD18 blocking antibody resulted in a more pronounced increase in γδ IEL migratory speed relative to anti-CD11c blockade. These data indicate that CD47 may regulate the affinity of CD18 heterodimers to reduce the adhesive interactions^24^ required for efficient γδ IEL migration. Taken together, our findings suggest that CD47 activation of CD11a/CD18 may be able to compensate for the loss of CD11c/CD18 binding. However, disruption of CD11c/CD18 in conjunction with impaired CD47-mediated signaling, either through CD11a/CD18 or other ligands, may reduce γδ IEL adhesion dynamics resulting in faster surveillance.

In the context of T cell motility, CD11a/CD18 (lymphocyte function-associated antigen-, LFA-1) and CD11c/CD18 can bind to Intercellular Adhesion Molecule-1 (ICAM-1)^25,26^. ICAM-1 is upregulated on the apical surface of intestinal epithelial cells in response to inflammation to facilitate CD11b/CD18-mediated neutrophil TEM into the lumen^7,27^, but ICAM-1 expression along the basolateral aspect of the epithelium has not been described. Epithelial occludin is concentrated at the tight junction but also diffuses within the basolateral membrane to promote γδ IEL motility^3,28^, presenting the possibility that low levels of basolateral ICAM-1 may modulate γδ IEL retention within the LIS via CD47-induced CD11a/CD18 signaling. Alternatively, CD11c/CD18 binds to fibrinogen^29^, a component of the basement membrane constitutively secreted by intestinal epithelial cells^30^. The loss of CD11c/CD18-mediated binding to this matrix protein could compound the adhesion defect caused by loss of CD47 signaling which may explain the increased γδ IEL migratory speed observed in CD47 KO^γδ^ mice. We cannot rule out the possibility that additional epithelial binding partners for this integrin complex and/or CD47 have yet to be identified.

We were intrigued by our observation of increased γδ IEL motility and reduced dwell time in the LIS in CD47 KO^IEC^ mice since several aspects of this migratory phenotype overlap with CD103 KO γδ IELs. We previously reported that CD103-deficient mice are less susceptible to bacterial invasion, as the enhanced surveillance of γδ IELs in these mice prevented microbial invasion^3,5^. To our surprise, the altered migratory behavior in CD47 KO^IEC^ mice had the opposite effect, with a trend toward increased *Salmonella* translocation despite an overall enhancement of γδ IEL migratory speed. These results indicate that the frequency of γδ IELs in the LIS is likely the determining factor contributing to γδ IEL-mediated protection against luminal pathogens, highlighting the need for further investigation into the underlying mechanisms regulating γδ IEL surveillance behavior in response to infection and inflammation.

In summary, we establish complementary yet distinct roles for epithelial and γδ T cell CD47 in regulating γδ IEL migration in the murine small intestine at steady-state. Further, we begin to elucidate the integrin-mediated mechanism by which CD47 influences γδ IEL motility, identifying CD18 as an additional regulator of homeostatic γδ IEL migration. A recent report highlights CD11c as one of many markers indicative of an effector-like γδ IEL phenotype^31^, suggesting that γδ IEL differentiation state may also influence surveillance behavior. This finding is notable as we recently reported that γδ IEL motility is severely compromised prior to the onset of Crohn’s disease-like ileitis which was concomitant with the influx of immature γδ T cells into the epithelial compartment^32^. Although the cause of impaired γδ IEL motility during preclinical inflammation remains elusive, we posit that the inability to upregulate key integrins such as CD11c may influence their surveillance behavior. The absence of commensal microbes leads to an increased proportion of stem-like γδ IELs^31^, and migration into the LIS is reduced in gnotobiotic and antibiotic-treated mice^33,34^. While these observations are correlative, our findings suggest that surface molecules induced during γδ IEL differentiation may contribute to their surveillance behavior. Failure to upregulate specific integrins or other epithelial binding partners may compromise barrier surveillance to increase the likelihood of microbial translocation and disease initiation.

## Acknowledgments

Research reported in this publication was supported by the Flow Cytometry CoRE (RRID:SCR_027701) at the Icahn School of Medicine at Mount Sinai and by the National Cancer Institute of the National Institutes of Health under award number P30 CA196521. The content is solely the responsibility of the authors and does not necessarily represent the official views of the National Institutes of Health. Microscopy and/or image analysis was performed at the Microscopy and Advanced Bioimaging CoRE at the Icahn School of Medicine at Mount Sinai.

## Author contributions

A.P. designed and performed experiments and wrote the manuscript. M.A.F. designed and performed experiments and analyzed the data. C.P. provided resources and advised the research. K.L.E. conceived the study, supervised the research, analyzed the data and wrote the manuscript. All authors approved the final manuscript.

## Supplementary Figures

**Supplementary Figure 1:**
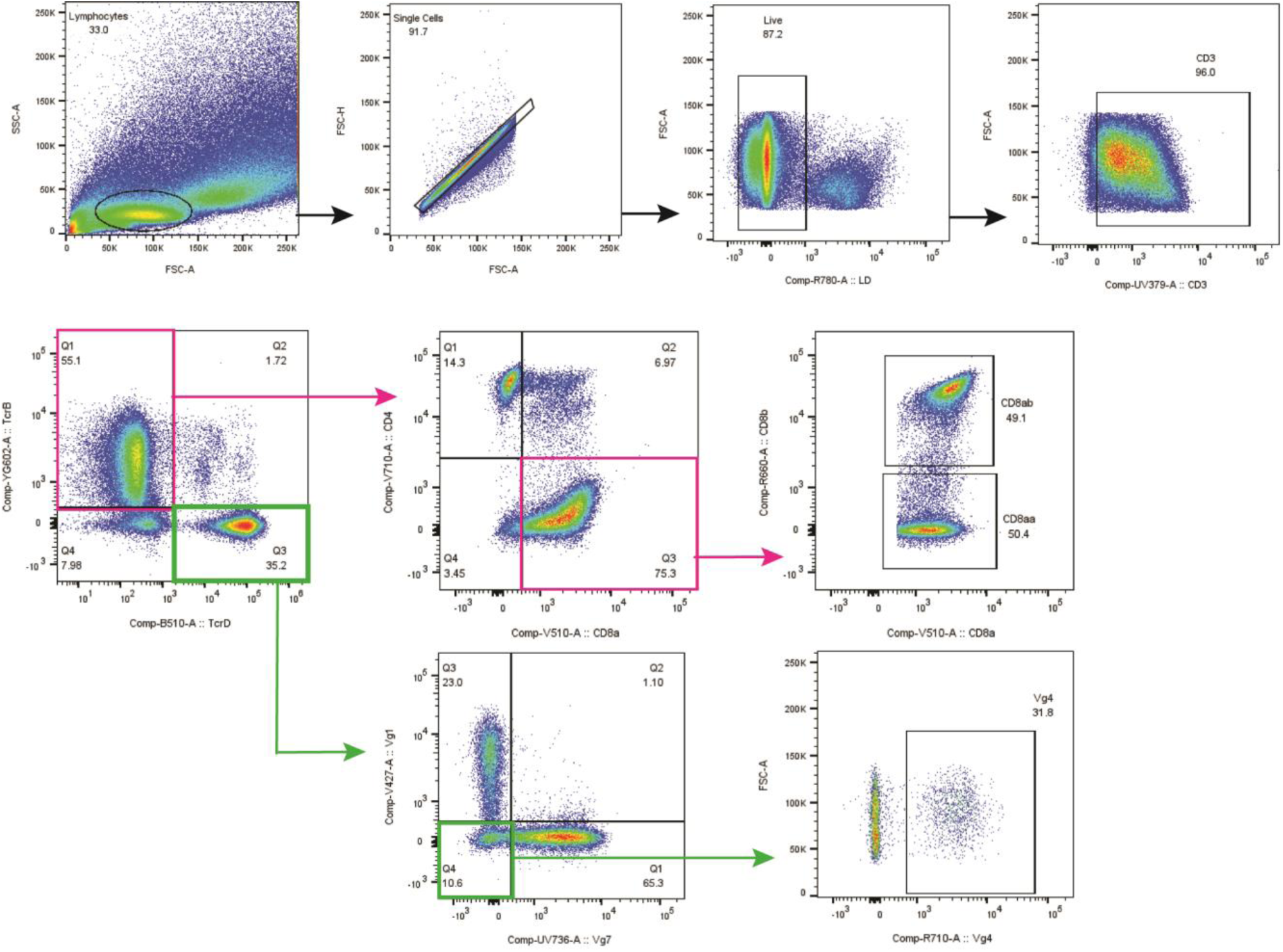
Gating strategy for intraepithelial lymphocyte subsets. IELs were isolated from the small intestine and CD3^+^ T cells were gated off of live single cells followed by TCRγδ or TCRβ. TCRβ* IEL subsets were subsequently gated based on CD4 and/or CD8α. and then CD8αβ or CD8αα. Conversely, γδ IELs were gated by Vγ subset.

**Supplementary Figure 2:**
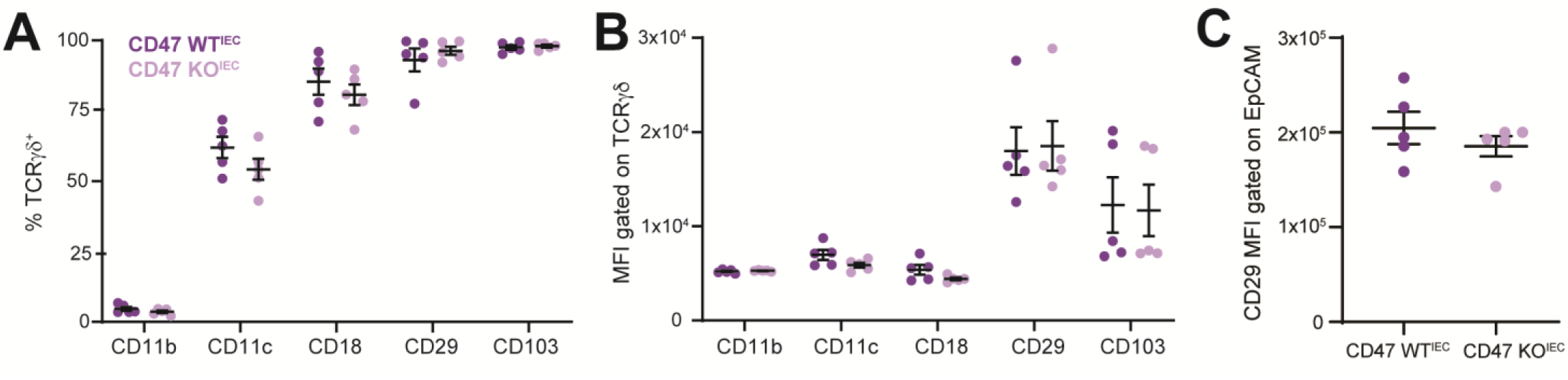
lntegrin expression is not altered in mice with CD47-deficient enterocytes. (A) Frequency and (B) MFI of the expression of various integrins on γδ IELs or (C) MFI of CD29 (β1 integrin) on EpCAM^+^ epithelial cells isolated from CD47 WT^IEC^ or CD47 KO^IEC^ mice. Mean ± SEM. Data shown from two independent experiments. Each data point represents an individual mouse. n=5. Statistical analysis: (A,B) two-way ANOVA with Sidak’s posthoc test, (C) Mann-Whitney U test.

**Supplementary Video 1**. Intravital microscopy of γδ T cells (green), luminal Alexa Fluor 633 (magenta) and nuclei (white) in jejunum of CD47 het^γδ^ (left) or CD47 KO^γδ^ (right) mice. Frames were collected approximately every 10 s. Tracks shown reflect the last 150 s of migration.

**Supplementary Video 2**. Intravital microscopy of γδ T cells (green), luminal Alexa Fluor 633 (magenta) and nuclei (white) in jejunum of CD47 het^γδ^ (left) or CD47 KO^γδ^ (right) mice following treatment with anti-CD11c. Frames were collected approximately every 10 s. Tracks shown reflect the last 150 s of migration.

**Supplementary Video 3**. Intravital microscopy of γδ T cells (green), luminal Alexa Fluor 633 (magenta) and nuclei (white) in jejunum of CD47 het^γδ^ (left) or CD47 KO^γδ^ (right) mice following treatment with anti-CD18. Frames were collected approximately every 10 s. Tracks shown reflect the last 150 s of migration.

## Abbreviations

EpCAM: epithelial cell adhesion molecule
GFP: green fluorescent protein
IAP: integrin associated protein
ICAM-1: Intercellular Adhesion Molecule 1
IEL: intraepithelial lymphocyte
IFNγ: interferon-γ
KO: knockout
LFA-1: lymphocyte function-associated antigen-1
LIS: lateral intercellular space
SIRPα: signal-regulatory protein alpha
TCR: T cell receptor
TEM: transepithelial migration
TSP-1: thrombospondin-1
WT: wildtype

## References

1 Hu, M. D. & Edelblum, K. L. Sentinels at the frontline: the role of intraepithelial lymphocytes in inflammatory bowel disease. Curr Pharmacol Rep 3, 321–334 (2017). 10.1007/s40495-017-0105-2

2 Cheroutre, H., Lambolez, F. & Mucida, D. The light and dark sides of intestinal intraepithelial lymphocytes. Nat Rev Immunol 11, 445–456 (2011). 10.1038/nri3007

3 Edelblum, K. L. et al. Dynamic migration of gammadelta intraepithelial lymphocytes requires occludin. Proc Natl Acad Sci U S A 109, 7097–7102 (2012). 10.1073/pnas.1112519109

4 Hu, M. D., Jia, L. & Edelblum, K. L. Policing the intestinal epithelial barrier: Innate immune functions of intraepithelial lymphocytes. Curr Pathobiol Rep 6, 35–46 (2018).

5 Edelblum, K. L. et al. gammadelta Intraepithelial Lymphocyte Migration Limits Transepithelial Pathogen Invasion and Systemic Disease in Mice. Gastroenterology 148, 1417–1426 (2015). 10.1053/j.gastro.2015.02.053

6 Brazil, J. C. & Parkos, C. A. Pathobiology of neutrophil-epithelial interactions. Immunol Rev 273, 94–111 (2016). 10.1111/imr.12446

7 Parkos, C. A. et al. CD47 mediates post-adhesive events required for neutrophil migration across polarized intestinal epithelia. J Cell Biol 132, 437–450 (1996).

8 Liu, Y. et al. The role of CD47 in neutrophil transmigration. Increased rate of migration correlates with increased cell surface expression of CD47. J Biol Chem 276, 40156–40166 (2001). 10.1074/jbc.M104138200

9 Azcutia, V. et al. Neutrophil expressed CD47 regulates CD11b/CD18-dependent neutrophil transepithelial migration in the intestine in vivo. Mucosal Immunol 14, 331–341 (2021). 10.1038/s41385-020-0316-4

10 Brown, E. J. & Frazier, W. A. Integrin-associated protein (CD47) and its ligands. Trends Cell Biol 11, 130–135 (2001).

11 Liu, Y. et al. Signal regulatory protein (SIRPalpha), a cellular ligand for CD47, regulates neutrophil transmigration. J Biol Chem 277, 10028–10036 (2002). 10.1074/jbc.M109720200

12 Liu, Y. et al. Peptide-mediated inhibition of neutrophil transmigration by blocking CD47 interactions with signal regulatory protein alpha. J Immunol 172, 2578–2585 (2004). 10.4049/jimmunol.172.4.2578

13 Lindberg, F. P. et al. Decreased resistance to bacterial infection and granulocyte defects in IAP-deficient mice. Science 274, 795–798 (1996).

14 Brenes, A. J. et al. Tissue environment, not ontogeny, defines murine intestinal intraepithelial T lymphocytes. Elife 10 (2021). 10.7554/eLife.70055

15 Reed, M. et al. Epithelial CD47 is critical for mucosal repair in the murine intestine in vivo. Nat Commun 10, 5004 (2019). 10.1038/s41467-019-12968-y

16 Jia, L. & Edelblum, K. L. Intravital Imaging of Intraepithelial Lymphocytes in Murine Small Intestine. J Vis Exp (2019). 10.3791/59853

17 Willingham, S. B. et al. The CD47-signal regulatory protein alpha (SIRPa) interaction is a therapeutic target for human solid tumors. Proc Natl Acad Sci U S A 109, 6662–6667 (2012). 10.1073/pnas.1121623109

18 Blazar, B. R. et al. Cd47 (Integrin-Associated Protein) Engagement of Dendritic Cell and Macrophage Counterreceptors Is Required to Prevent the Clearance of Donor Lymphohematopoietic Cells. Journal of Experimental Medicine 194, 541–550 (2001). 10.1084/jem.194.4.541

19 Barclay, A. N. Signal regulatory protein alpha (SIRPalpha)/CD47 interaction and function. Curr Opin Immunol 21, 47–52 (2009). 10.1016/j.coi.2009.01.008

20 Oldenborg, P.-A. et al. Role of CD47 as a Marker of Self on Red Blood Cells. Science 288, 2051–2054 (2000). doi:10.1126/science.288.5473.2051

21 Komori, S. et al. CD47 promotes peripheral T cell survival by preventing dendritic cell-mediated T cell necroptosis. Proc Natl Acad Sci U S A 120, e2304943120 (2023). 10.1073/pnas.2304943120

22 Huleatt, J. W. & Lefrancois, L. Antigen-driven induction of CD11c on intestinal intraepithelial lymphocytes and CD8+ T cells in vivo. J Immunol 154, 5684–5693 (1995).

23 McIntyre, C. L. et al. beta2 Integrins differentially regulate gammadelta T cell subset thymic development and peripheral maintenance. Proc Natl Acad Sci U S A 117, 22367–22377 (2020). 10.1073/pnas.1921930117

24 Azcutia, V. et al. CD47 plays a critical role in T-cell recruitment by regulation of LFA-1 and VLA-4 integrin adhesive functions. Mol Biol Cell 24, 3358–3368 (2013). 10.1091/mbc.E13-01-0063

25 Edwards, C. P. et al. Identification of amino acids in the CD11a I-domain important for binding of the leukocyte function-associated antigen-1 (LFA-1) to intercellular adhesion molecule-1 (ICAM-1). J Biol Chem 270, 12635–12640 (1995). 10.1074/jbc.270.21.12635

26 Blackford, J., Reid, H. W., Pappin, D. J., Bowers, F. S. & Wilkinson, J. M. A monoclonal antibody, 3/22, to rabbit CD11c which induces homotypic T cell aggregation: evidence that ICAM-1 is a ligand for CD11c/CD18. Eur J Immunol 26, 525–531 (1996). 10.1002/eji.1830260304

27 Sumagin, R., Robin, A. Z., Nusrat, A. & Parkos, C. A. Transmigrated neutrophils in the intestinal lumen engage ICAM-1 to regulate the epithelial barrier and neutrophil recruitment. Mucosal Immunol 7, 905–915 (2014). 10.1038/mi.2013.106

28 Shen, L., Weber, C. R. & Turner, J. R. The tight junction protein complex undergoes rapid and continuous molecular remodeling at steady state. J Cell Biol 181, 683–695 (2008). 10.1083/jcb.200711165

29 Loike, J. D. et al. CD11c/CD18 on neutrophils recognizes a domain at the N terminus of the A alpha chain of fibrinogen. Proc Natl Acad Sci U S A 88, 1044–1048 (1991). 10.1073/pnas.88.3.1044

30 Seltana, A. et al. Fibrin(ogen) Is Constitutively Expressed by Differentiated Intestinal Epithelial Cells and Mediates Wound Healing. Front Immunol 13, 916187 (2022). 10.3389/fimmu.2022.916187

31 Xie, S. et al. Effector differentiation by stem-like intraepithelial gammadelta T cells is required for host defense against infection. Immunity 59, 577–597 e577 (2026). 10.1016/j.immuni.2026.01.006

32 Xu, W. et al. Dysregulation of gammadelta intraepithelial lymphocytes precedes Crohn’s disease-like ileitis. Sci Immunol 10, eadk7429 (2025). 10.1126/sciimmunol.adk7429

33 Jia, L. et al. A transmissible gammadelta intraepithelial lymphocyte hyperproliferative phenotype is associated with the intestinal microbiota and confers protection against acute infection. Mucosal Immunol 15, 772–782 (2022). 10.1038/s41385-022-00522-x

34 Hoytema van Konijnenburg, D. P., et al. Intestinal Epithelial and Intraepithelial T Cell Crosstalk Mediates a Dynamic Response to Infection. Cell 171, 783–794 (2017). 10.1016/j.cell.2017.08.046

